# Dynamic cortical routing mediates temporal attention

**DOI:** 10.64898/2026.04.03.716160

**Authors:** Jiating Zhu, Karen J. Tian, Marisa Carrasco, Rachel N. Denison

## Abstract

Selecting information from dynamic streams requires mechanisms that prioritize a visual stimulus at a specific moment over preceding and subsequent stimuli at the same location. Whereas selective temporal attention has been found to enhance neural responses to stimuli, its impact on communication between brain regions remains unexplored. Here, we investigated whether prioritizing a stimulus at a specific time is achieved through selective routing of stimulus information across cortical networks using MEG. We developed a dynamic informational connectivity approach to quantify shared stimulus information between each region and the rest of the network. When stimuli compete in time, we found that temporal attention modulated the network at both early and late post-target time windows, routing information along two possible pathways—occipito-fronto-cingulate and occipito-temporal—via both transient bursts of network communication and theta-rhythmic replay. These results provide evidence that under dynamic sensory input, the timing of neural communication determines stimulus selection.

## Introduction

Sensory information is continuous and dynamic, yet the capacity to process such information is limited. To manage these constraints, humans use attention to prioritize information most relevant to their current task. For example, when caring for a baby, it is critical to prioritize parts of the visual stream that signal imminent risk—such as the baby’s hand moving toward a small object. Flexible attentional prioritization enables relevant information to guide goal-directed behavior, like moving the object out of the baby’s reach (Carrasco, 2011; Anton-Erxleben and Carrasco, 2013; Nobre and Van Ede, 2023; Denison, 2024).

Attention is thought to prioritize task-relevant information via selective routing: the preferential communication across cortical networks of some stimulus information at the expense of other information. Attention enhances sensory responses to task-relevant stimuli (van Es et al., 2018; Dugué et al., 2020; Liu et al., 2021; Maunsell and Treue, 2006; Liu et al., 2007; Foster and Ling, 2022), potentially increasing the likelihood that the associated information is transmitted downstream. Attention also directly increases inter-areal communication by synchronizing neuronal activity across areas, such that spikes in an upstream area are more effectively transmitted to a downstream area (Fries, 2015; Salinas and Sejnowski, 2001). However, our understanding of selective routing remains limited in several respects. First, selective routing has mainly been inferred from synchronized neuronal oscillations (Banaie Boroujeni et al., 2025; Fries, 2015; Saalmann et al., 2012; Womelsdorf et al., 2007), rather than by directly measuring the transmission of stimulus-specific information content (Van Ede et al., 2018; Kok et al., 2012; Stokes et al., 2015). Second, attentional modulation of inter-areal communication has largely been examined in selected pairs of brain regions (Fries, 2015; Gregoriou et al., 2009; Buschman and Miller, 2007; Saalmann et al., 2007), so little is known about how attention modulates routing across large-scale cortical networks (Banaie Boroujeni et al., 2025; Siegel et al., 2015). Third, selective routing has mainly been studied in the context of spatial attention (Fries, 2015; Bosman et al., 2012; Gregoriou et al., 2009; Buschman and Miller, 2007; Saalmann et al., 2007), whereas how temporal attention affects selective routing has not been investigated. Fourth, how selective routing unfolds over time for dynamic input has rarely been studied. Here we addressed these four limitations to advance understanding of how attention facilitates selective routing in dynamic settings.

Temporal attention refers to the prioritization of sensory information at specific points in time, which improves perception and behavior at attended moments (Denison et al., 2017, 2021; Fernández et al., 2019; Duyar et al., 2023, 2024; Palmieri and Carrasco, 2024; Denison et al., 2024; Tian et al., 2026; Jing et al., 2023; Huang et al., 2025). For example, returning to the caregiving scenario, by anticipating when the baby might grasp a dangerous object, the caregiver can attend at the critical moment to monitor the baby’s hand position and intervene if necessary. Attending too early or too late would make it difficult to respond appropriately. Temporal orienting increases neural responses to stimuli appearing at attended moments (Correa et al., 2006; Doherty et al., 2005), yet it poses a challenge for cortical routing mechanisms identified for spatial attention, which prioritizes the selective processing of one location over others presented at the same time. Spatial routing mechanisms exploit spatiotopic organization by facilitating communication along spatially specific routes, in effect selecting one of multiple routes for preferential transmission. Temporal attention, however, can prioritize stimuli presented at the same spatial location but at different moments. Selective routing in the temporal domain therefore cannot rely on separate cortical populations or communication routes. Instead, we hypothesized that temporal attention flexibly shapes the dynamics of selective routing over a shared network pathway.

To investigate whether and how temporal attention alters selective routing across large-scale cortical networks, we developed an analysis approach called “dynamic informational connectivity.” This approach is based on the informational connectivity method, which quantifies correlated fluctuations in multivariate pattern discriminability across regions (Coutanche and Thompson-Schill, 2013), thereby directly measuring shared fluctuations in stimulus-specific content across brain regions. Whereas previous studies have typically calculated a single connectivity estimate from extended functional magnetic resonance imaging (fMRI) time series (Coutanche and Thompson-Schill, 2013; Anzellotti and Coutanche, 2018; Huang et al., 2024; Mill and Cole, 2025), here we leveraged the high temporal resolution of magnetoencephalography (MEG) and sliding time windows to capture rapid fluctuations in informational connectivity. To quantify informational connectivity across large-scale networks, we combined this approach with graph-theory metrics.

We recorded MEG while human participants performed a temporal attention task, in which they were instructed to attend to one of two consecutive target stimuli presented at the same spatial location, with one target more likely to be task relevant. We used source-reconstruction to estimate neural activity across large-scale cortical networks during this task. To track the flow of stimulus information—and to overcome the challenge that successive stimuli may engage overlapping neural populations—we independently decoded the identity of each stimulus and quantified how decoding accuracy, as a measure of stimulus information, was shared across areas and networks in a time-resolved fashion.

Our results revealed that temporal attention alters informational connectivity across large-scale cortical networks. Temporal attention led to discrete, stimulus-specific bursts of increased connectivity at both early and late time windows during the trial, indicating that selective routing is neither instantaneous nor sustained continuously. The connectivity changes coalesced along two principal pathways—an occipital-temporal route and an occipital-fronto-cingulate route—which appeared to play different roles in maintaining stimulus information throughout the trial and mediating the transition from processing the first to the second target, respectively. Finally, we observed a periodic recurrence at 4 Hz of occipital decoding patterns within the temporal lobe following both targets. Altogether, these findings reveal how temporal attention dynamically and selectively routes stimulus information across large-scale cortical networks in the service of flexible behavior.

## Results

### Temporal attention improved orientation discrimination

Participants performed a two-target temporal cueing task while MEG was recorded (**Figure 1A**). Each trial presented two sequential gratings separated by a 300 ms stimulus onset asynchrony. A precue tone at the start of the trial instructed participants to attend to either the first (T1) or the second (T2) target, and a response cue at the end of the trial indicated which target’s tilt should be reported. Participants reported whether the tilt of the indicated target was clockwise or counterclockwise relative to the vertical or horizontal axis (**Figure 1B**). On 75% of trials the precue and response cue matched (valid) and on 25% they mismatched (invalid), incentivizing participants to prioritize the precued target. Gratings were tilted independently around either the vertical or horizontal axis, enabling MEG decoding of axis orientation—a feature orthogonal to the participant’s report. Temporal attention improved tilt discriminability (**Figure 1C**; main effect of validity: *F* (1, 9) = 20.22, p = 0.0015,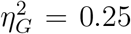) and response times (**Figure 1D**; main effect of validity: *F* (1, 9) = 70.60, p *<* 0.001,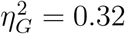) for both targets. Full behavioral results were reported in (Zhu et al., 2024); see also **Supplementary Figure 1**.

**Figure 1.**
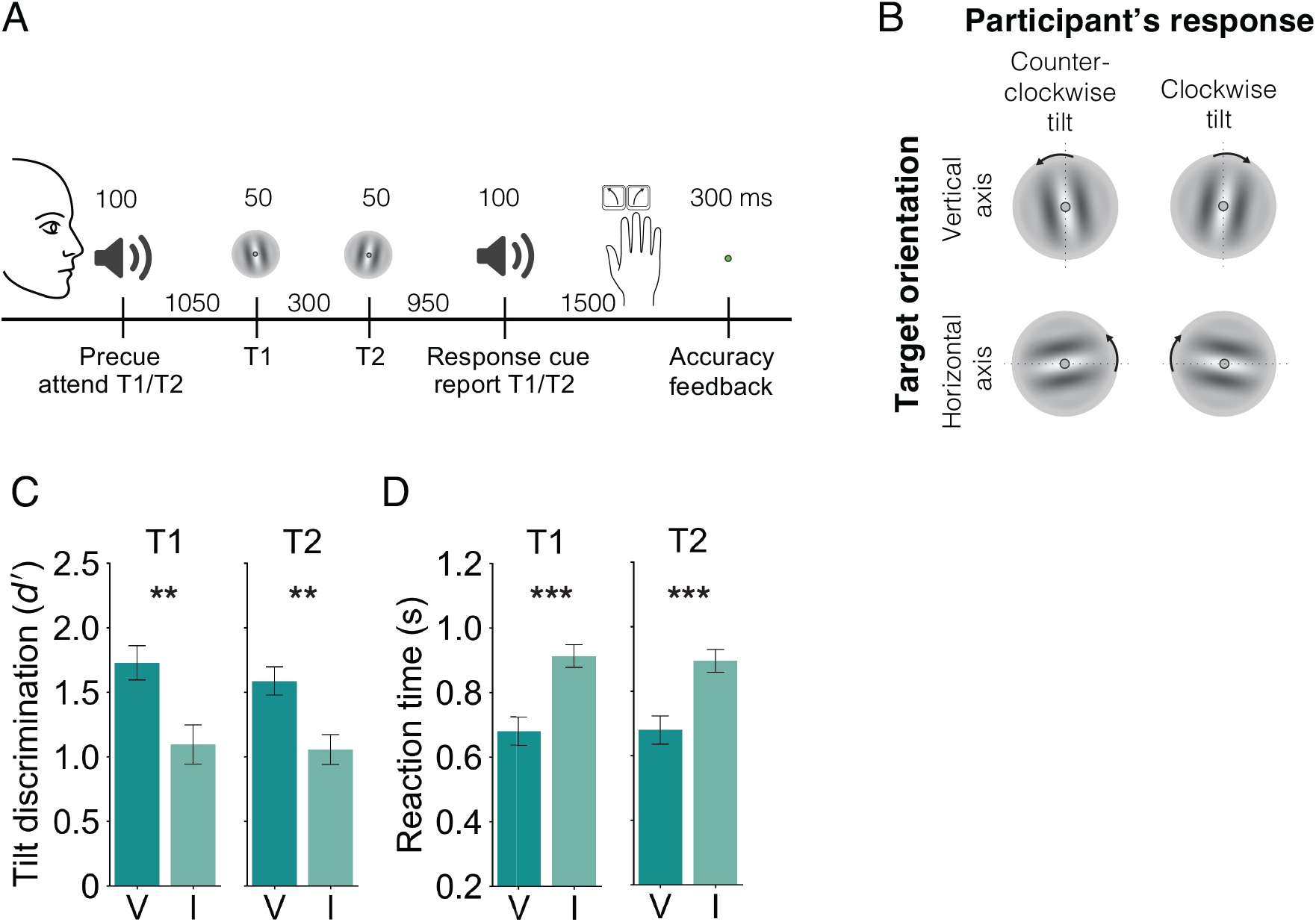
Two-target temporal cueing task. **(A)** Trial timeline showing stimulus durations and SOAs. Precues and response cues were pure tones (high = cue T1, low = cue T2). **(B)** Each target was independently tilted along either the vertical or horizontal axis. Participants reported whether the target specified by the response cue was tilted clockwise or counterclockwise, while the axis orientation served as the stimulus feature decoded in the analysis. **(C)** Tilt discrimination (sensitivity) and **(D)** reaction time for each target (T1, T2) by validity condition. Results are previously reported in (Zhu et al., 2024). Sensitivity was higher and reaction time was faster for valid (V) than invalid (I) trials. Error bars indicate ±1 SEM. ** p *<* 0.01; *** p *<* 0.001.

### Dynamic informational connectivity analysis

To investigate the flow of stimulus information across the cortex, we used informational connectivity (Coutanche and Thompson-Schill, 2013; Anzellotti and Coutanche, 2018) to measure shared fluctuations in stimulus information across brain regions. For each atlas-based brain region (34 per hemisphere, (Desikan et al., 2006), **Supplementary Table 1**) from MEG source-reconstructed data, and for each target separately, we decoded stimulus orientation (vertical vs. horizontal) from the multivariate neural activity pattern with 5-ms resolution. Then we calculated the correlation between the decoding accuracy time series from each pair of regions in the cortex. We used a 50-ms sliding time window to estimate informational connectivity in a time-resolved fashion across the trial, yielding a dynamic informational connectivity analysis (**Figure 2**). This procedure, applied to all pairs of brain regions, produced an informational connectivity network for each target (T1 or T2), time window, and attention condition (target attended or unattended). For subsequent graph theory analyses on these networks, each atlas-based region was considered a node, and the correlation in stimulus orientation decoding accuracy for a pair of regions was considered an edge.

**Figure 2.**
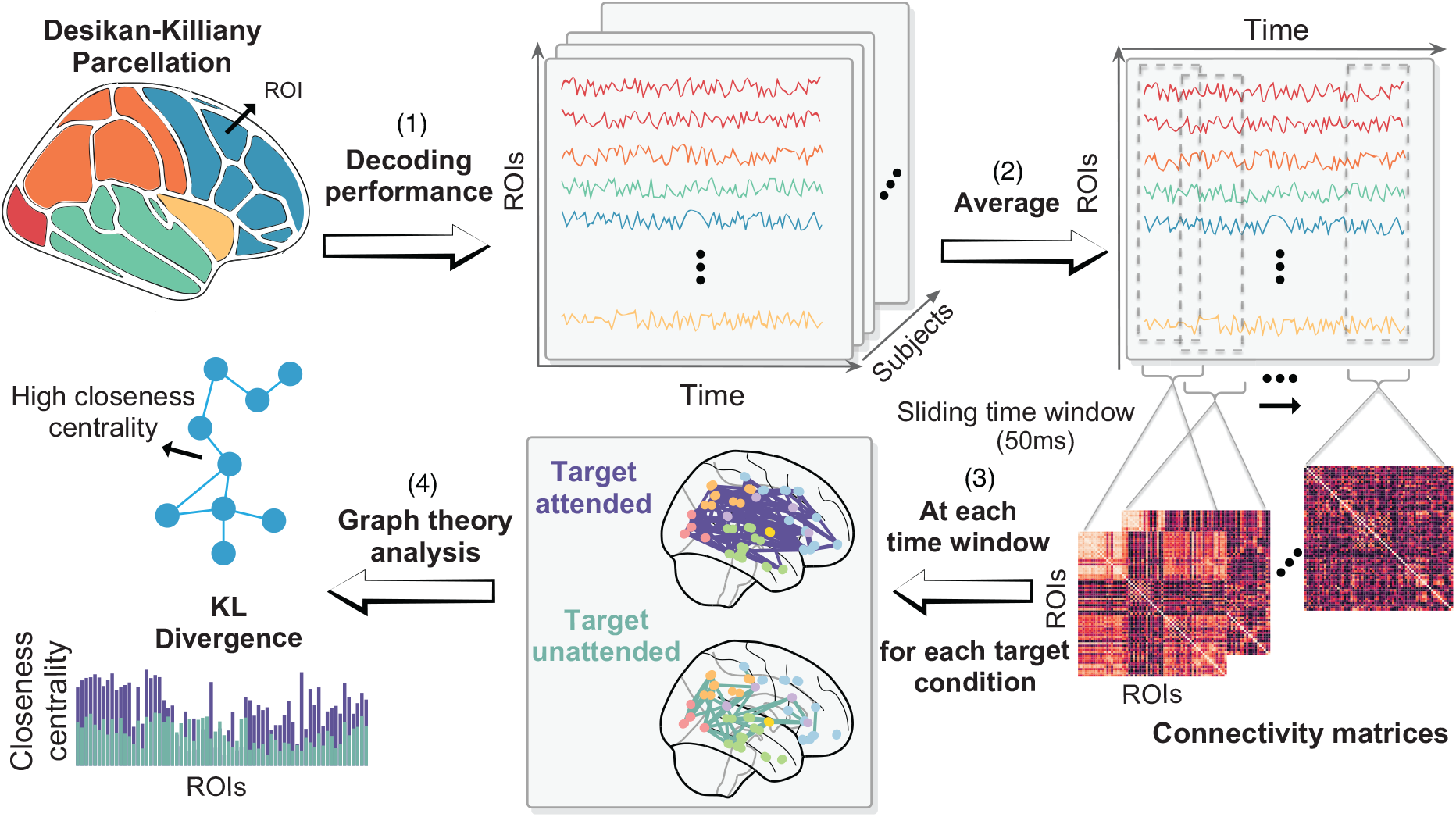
Whole-brain comparison of informational connectivity between target-attended and target-unattended conditions. (1) For each region, we trained an SVM classifier to decode target axis orientation from source-reconstructed MEG data in a time-resolved fashion. (2) Decoding accuracy for each region was averaged across subjects. (3) For each target condition, we then computed the correlation of decoding accuracy (edges) across all pairs of atlas-based regions (nodes) within sliding time windows. (4) At each time window, closeness centrality for each region was defined as the average inverse correlation distance to all other regions. This yielded a distribution of closeness centrality across all regions. Finally, we compared the spatial patterns of closeness centrality between conditions using Kullback–Leibler (KL) divergence.

We first aimed to determine whether temporal attention altered the informational connectivity network structure across time. To do so, we estimated the degree of stimulus information sharing, separately for each target, between each brain region and the rest of the network at each time step. We quantified information sharing using the graph theory property of closeness centrality, calculated as the average inverse correlation distance from one node to all other nodes. Regions with higher closeness centrality in a given time window thus exhibit more similar orientation decoding time series to other brain regions within that time window. To assess effects of attention on the network structure, we used Kullback-Leibler (KL) divergence to compare closeness centrality distributions (across nodes) for target attended vs. unattended networks, again for each target and time window (see **Methods**, *Dynamic informational connectivity analysis*). This procedure provided a data-driven approach to assess whether attention to a target changed how stimulus information for that target was shared across the cortical network over time. This approach goes beyond assessing correlations between specific pairs of regions by summarizing the network structure of informational connectivity.

### Temporal attention changed informational connectivity at both early and late post-target time windows

#### T1 informational network

The KL divergence between T1 attended and unattended networks (**Figure 3A**) showed that temporal attention changed the spatial pattern of closeness centrality in three time windows: early (50-130 ms after T1 onset, p = 0.04 cluster corrected one-sided permutation test); intermediate (250-335 ms after T1 onset p = 0.03 cluster corrected), right before T2 onset; and late, 625-730 ms after T1 onset (~ 300 ms after T2 onset, p *<* 0.001 cluster corrected). To investigate which brain areas contributed to these network differences, we obtained closeness centrality measures for the occipital, parietal, temporal, frontal, and cingulate lobes by averaging closeness centrality values across the atlas-based regions within each lobe within each significant time window.

**Figure 3.**
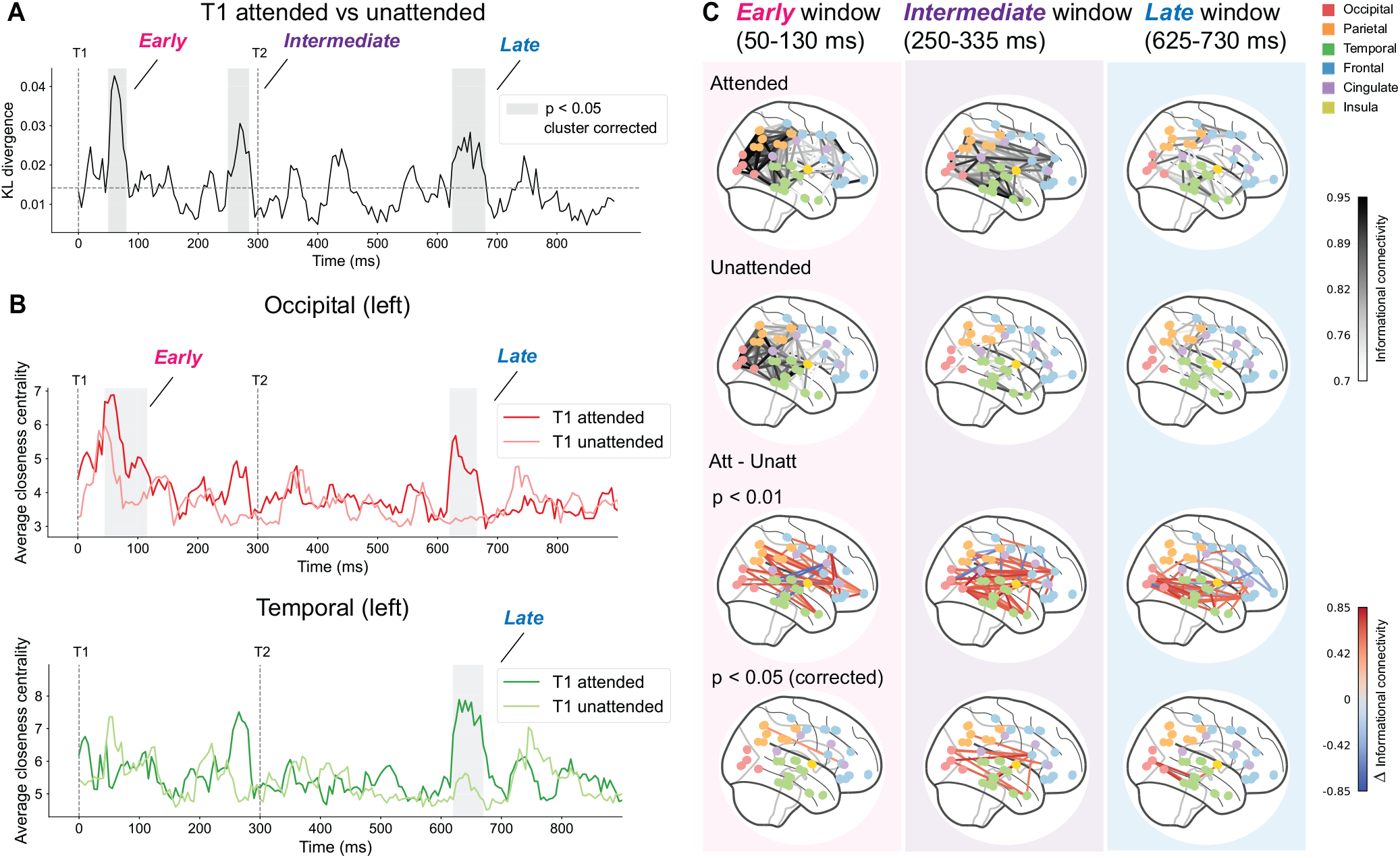
Modulation of informational connectivity for T1. Plotted time series values are indexed to the time point at the start of the sliding window used to compute the value. **(A)** KL divergence between T1 attended vs. T1 unattended informational connectivity networks indicate significant divergence at three time windows (early 50-130 ms, intermediate 250-335 ms, late 625-730 ms) from T1 onset. The horizontal dashed line marks the average value over the time window of interest (0-900 ms), as a baseline measure. Each shaded window shows the range between starting time points of the first and last sliding windows in the significant interval, whereas the range reported in text comprises the starting time point of first to the ending time point of the last sliding window in the interval. **(B)** Temporal attention enhanced average closeness centrality values for ROIs in the left occipital lobe at an early (45-165 ms) and a late (620-715 ms) time window after target onset, and in left temporal lobe in a late (620-720 ms) window. Gray shaded regions indicate cluster corrected p *<* 0.05 (threshold were 0.85 for the early window and 0.95 for the late windows). The same shading convention as in **(A). (C)** Average informational connectivity at the early, intermediate, and late time windows. The visualization of target attended networks is on the first row, and target unattended networks on the second row. Darker edges indicate stronger connectivity. The third row shows the difference in informational connectivity between attended and unattended conditions. The presented edges are significant at p *<* 0.01 compared to a shuffled null distribution, with red edges indicating higher attended connectivity. The fourth row presents the FDR corrected edges (p *<* 0.05).

We hypothesized that attention increases shared stimulus information across the network; thus, we expected higher closeness centrality for the attended than the unattended condition. Consistent with this prediction, we found that the average closeness centrality in the occipital lobe was enhanced in early (45-165 ms after T1 onset, p = 0.032 cluster corrected) and late (620-715 ms after T1 onset, p = 0.038 cluster corrected) time windows (**Figure 3B**). The average closeness centrality in the temporal lobe was also enhanced during the same late window (620-720 ms after T1 onset, p = 0.026 cluster corrected). These enhancements were specific to the left hemisphere. Other lobes did not exhibit significant closeness centrality changes with attention after cluster correction across the time series (p *>* 0.05).

Closeness centrality was calculated based on decoding accuracy, so we assessed whether the observed attentional increases in closeness centrality were accompanied by corresponding changes in decoding accuracy. Although decoding accuracy increased following the target, attention did not affect decoding accuracy in either left occipital or temporal lobe (**Supplementary Figure 2A**). These results indicate that the attentional changes in closeness centrality emerged at a network level, independent of local changes in decoding accuracy.

We next visualized the anatomical organization of informational connectivity across the network (**Figure 3C**). For each significant time window, we averaged the correlation values for each pair of nodes across all successive sliding windows (5 ms stride and 50 ms window size) contained within the significant window. Across all three time windows, the overall network connectivity was consistently stronger in the attended than the unattended condition. In the early time window, informational connectivity in the T1 attended and unattended networks was strongest in posterior brain areas. This pattern was expected given that stimulus information travels through the visual hierarchy starting from the occipital lobe. Occipital connectivity appeared stronger for the T1 attended than the T1 unattended network in the early time window (**Figure 3C, top two rows**), consistent with the closeness centrality time series analysis (**Figure 3B**). However, few individual connectivity edges survived FDR correction for multiple comparisons across edges in the early time window (**Figure 3C, bottom row**). In both the intermediate and late windows, there was a strong attentional enhancement of informational connectivity between nodes in the occipital and temporal lobes, which survived FDR correction (p *<* 0.05).

#### T2 informational network

Similar to T1, KL divergence between T2 attended and unattended networks (**Figure 4A**) showed that temporal attention changed the spatial pattern of closeness centrality at both early (“early-1”, 25-110 ms after T2 onset, p = 0.017 cluster corrected; “early-2”, 70-165 ms after T2 onset, p = 0.003 cluster corrected) and late time windows (450-535 ms after T2 onset, p = 0.046 cluster corrected). Closeness centrality averaged across the regions within each lobe showed temporal attentional enhancements in the early-1 time window in the occipital (20–95 ms after T2 onset, p = 0.035 cluster corrected) and temporal (20–100 ms after T2 onset, p = 0.013 cluster corrected) lobes, and the late window (405–525 ms after T2 onset, p = 0.045 cluster corrected) in the parietal lobe (**Figure 4C**).

**Figure 4.**
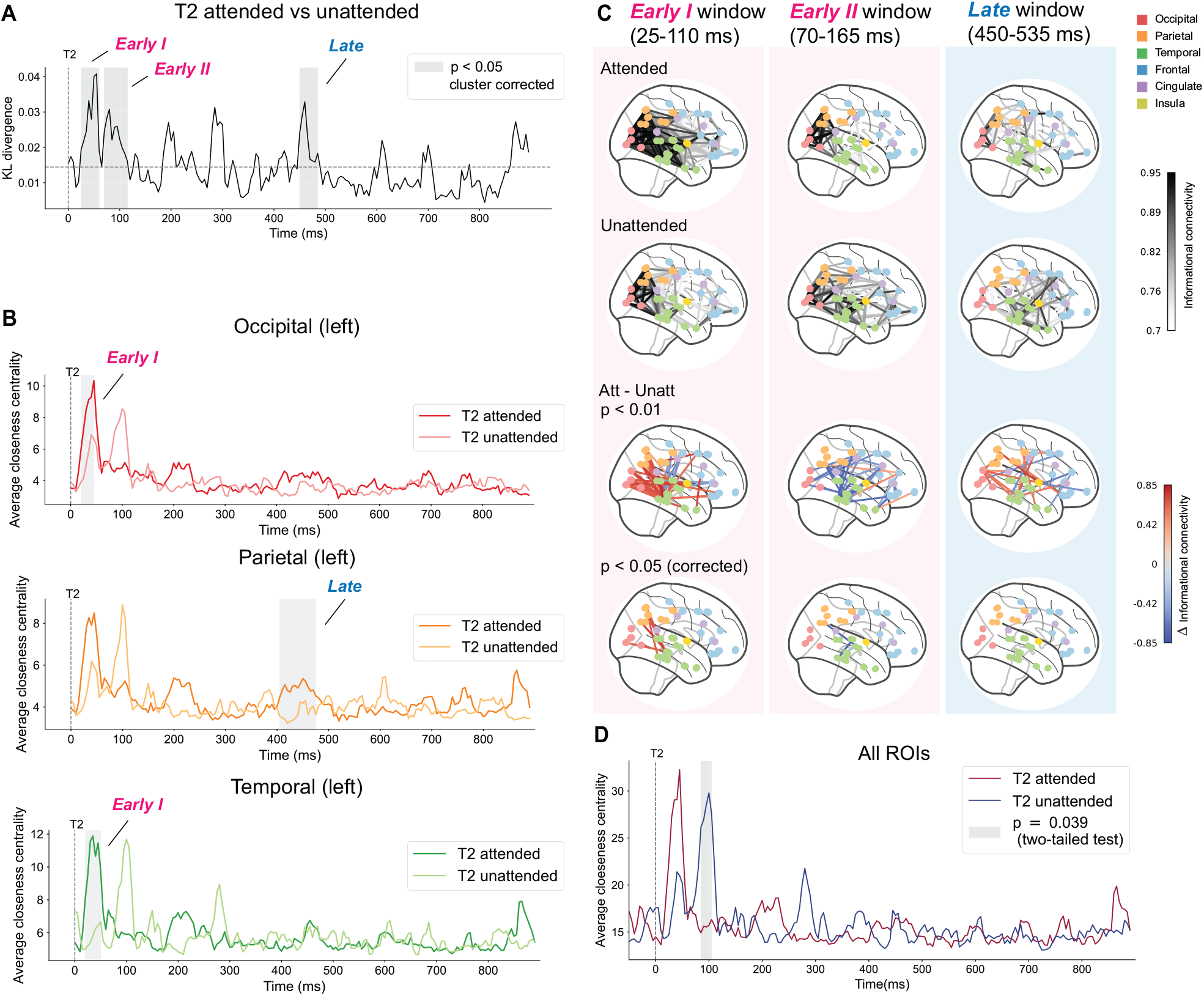
Modulation of informational connectivity for T2. **(A)** KL divergence between T2 attended vs. T2 unattended informational connectivity networks indicate significant divergence at early-1 (25-110 ms), early-2 (70-165 ms) and late (450-535 ms) time windows from T2 onset. The horizontal dashed line marks the average value over the time window of interest (0-900 ms). **(B)** Temporal attention enhanced average closeness centrality values for ROIs in the left occipital and temporal lobes at an early (20–95 ms) time window after target onset, and in the left parietal in a late (405–525 ms) window after target onset. Gray shaded regions indicate cluster corrected p *<* 0.05 (thresholds were 0.99 for the early windows and 0.85 for the late window). **(C)** Average informational connectivity at the early-1, early-2, and late time windows. The visualization of target attended networks is on the first row, and target unattended networks on the second row. Darker edges indicate stronger connectivity. The third row shows the difference in informational connectivity between attended and unattended conditions. The presented edges are significant at p *<* 0.01 compared to a shuffled null distribution, with red edges indicating higher attended connectivity. The fourth row presents the FDR corrected edges (p *<* 0.05). **(D)** Early modulation 85–155 ms after target onset for closeness centrality averaged all ROIs in the brain for T2. The peak for T2 unattended was delayed relative to the T2 attended peak. The threshold used for cluster correction here was 0.99. Plotting conventions are the same as for **Figure 3**.

When T2 was unattended, the increase in closeness centrality just after the target onset appeared to be delayed in a manner that was widespread across lobes. We evaluated this observation with a two-sided test on all ROIs and found that the closeness centrality of the T2 unattended network was significantly enhanced in the early-2 time window (85–155 ms after T2 onset, p = 0.039 cluster corrected two-sided permutation test), reflecting a delay of 60 ms in the early closeness centrality peak in unattended relative to attended trials (**Figure 4D**). The early-1 attentional enhancement, in contrast, was more localized in the left occipital and temporal lobes and did not survive across all lobes.

The attentional increases in closeness centrality for T2 were again not accompanied by attentional changes in decoding accuracy, with no significant decoding differences in the left occipital, left parietal, or left temporal lobes (**Supplementary Figure 2B**).

Visualization of the anatomical structure of informational connectivity (**Figure 4C**) confirmed increased connectivity in the attended condition compared to the unattended condition in the early-1 and late time windows. In contrast, T2 network connectivity was stronger in the unattended condition in the early-2 time window, consistent with the delayed closeness centrality peak shown in **Figure 4B** and **Figure 4D**. After FDR correction for multiple comparisons, the early-1 window showed significant connectivity differences between the occipital and temporal lobes as well as between the parietal and temporal lobes (p *<* 0.05).

In summary, both T1 and T2 showed attentional enhancements of informational connectivity between the occipital and temporal lobes, which occurred at intermediate and late time windows after T1 but at an early time window after T2. Meanwhile, T2 showed a striking delay in informational connectivity enhancement when T1 was attended, which did not occur for T1.

### Informational network dynamics during the transition from T1 to T2 processing

We next examined the anatomical organization of attention-modulated networks during the transition from T1 to T2 processing. When T1 was attended, the enhancement of T1 attended informational connectivity in the intermediate time window, right before T2, was left lateralized (**Figure 5, first column**). There were individual edges strongly connected (p *<* 0.01 uncorrected) the left rostral middle frontal, pars opercularis and posterior cingulate (see **Supplementary Figure 3**) regions in the frontal and cingulate lobes, which also showed enhanced decoding performance in the same time window (Zhu et al., 2024). After FDR correction for multiple comparisons, there were also significant connectivity differences between the left occipital and frontal lobes and between frontal and cingulate lobes (p *<* 0.05) in the T1 intermediate window.

**Figure 5.**
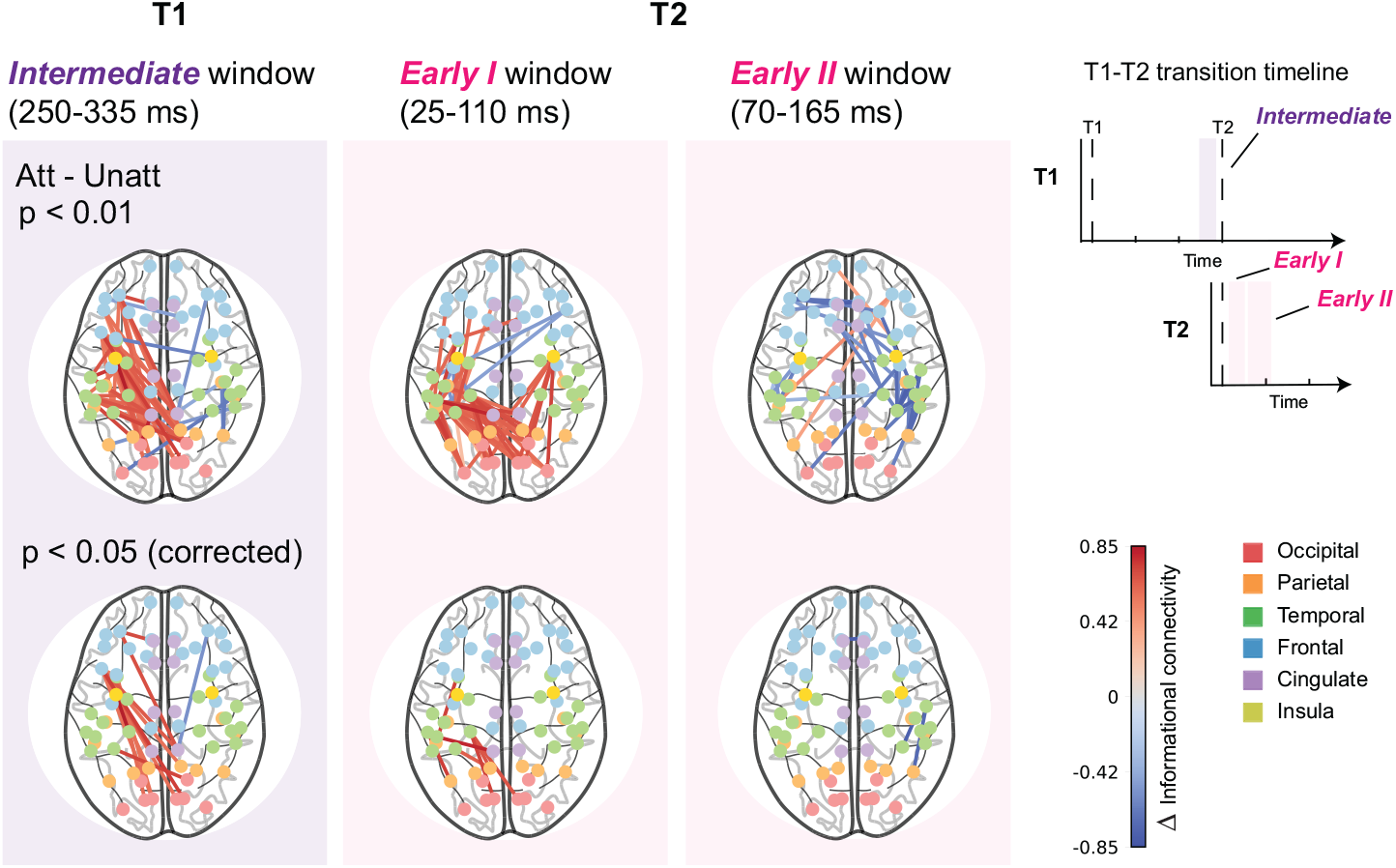
Axial view of the difference in informational connectivity between attended and unattended conditions during the transition from T1 to T2 processing. The first column shows the difference in T1 informational connectivity at the intermediate window (same data as in **Figure 3C**), occurring just before T2. The second and third columns show the difference in T2 informational connectivity at the early-1 and early-2 windows (same data as in **Figure 4C**), occurring just after T2. The presented edges in the first row are significant at p *<* 0.01 compared to a shuffled null distribution. Edges with higher values are shown in red, indicating greater connectivity during the attended than the unattended condition. The second row presents the FDR corrected edges (p *<* 0.05). The inset T1-T2 transition timeline shows the T1 processing timeline and T2 processing timeline over the same time axis.

When T2 was attended, attentional enhancements in the early-1 window, just after T2 onset, were similarly left lateralized (**Figure 5, second column**). After FDR correction for multiple comparisons, there were also significant connectivity differences between the temporal and frontal lobes (p *<* 0.05) in the T2 early-1 window. However, when T1 was attended and T2 was unattended, T2 informational connectivity was delayed (early-2 window) and right lateralized (**Figure 5, third column**). After FDR correction for multiple comparisons, there were significant connectivity differences between the parietal and temporal lobes (p *<* 0.05) in the T2 early-2 window.

Thus, just before and after the onset of T2, attention to T1 and to T2, respectively, increased informational connectivity along a left-lateralized pathway involving the occipital and frontal lobes.

### An early occipital motif recurred in the entorhinal-parahippocampal cortex

From the connectivity results above, we found that temporal attention enhanced informational connectivity between the occipital and temporal lobes for both targets. Temporal lobe regions subserving memory can be modulated by spatial attention (Wilming et al., 2018; Aly and Turk-Browne, 2016), and visual short-term memory plays a role in the current temporal attention task due to the requirement to wait for the response cue presented 1 s after T2 to know which stimulus to report. Therefore, we next asked how information was dynamically shared between the occipital lobe and the atlas-based regions within the temporal lobe most closely associated with memory, namely the entorhinal and parahippocampal cortex.

We asked how early visual information captured in the initial sensory response might be replayed in the entorhinal-parahippocampal cortex across time. To do so, we measured the relation between orientation information in the occipital lobe within the first 150 ms after target onset and information in the entorhinal–parahippocampal cortex across time by calculating the correlation between their decoding time series using sliding time windows (**Figure 6A**). For both T1 and T2, the correlation time series for the attended condition seemed to exhibit a periodic pattern (**Figure 6B, first row**).

**Figure 6.**
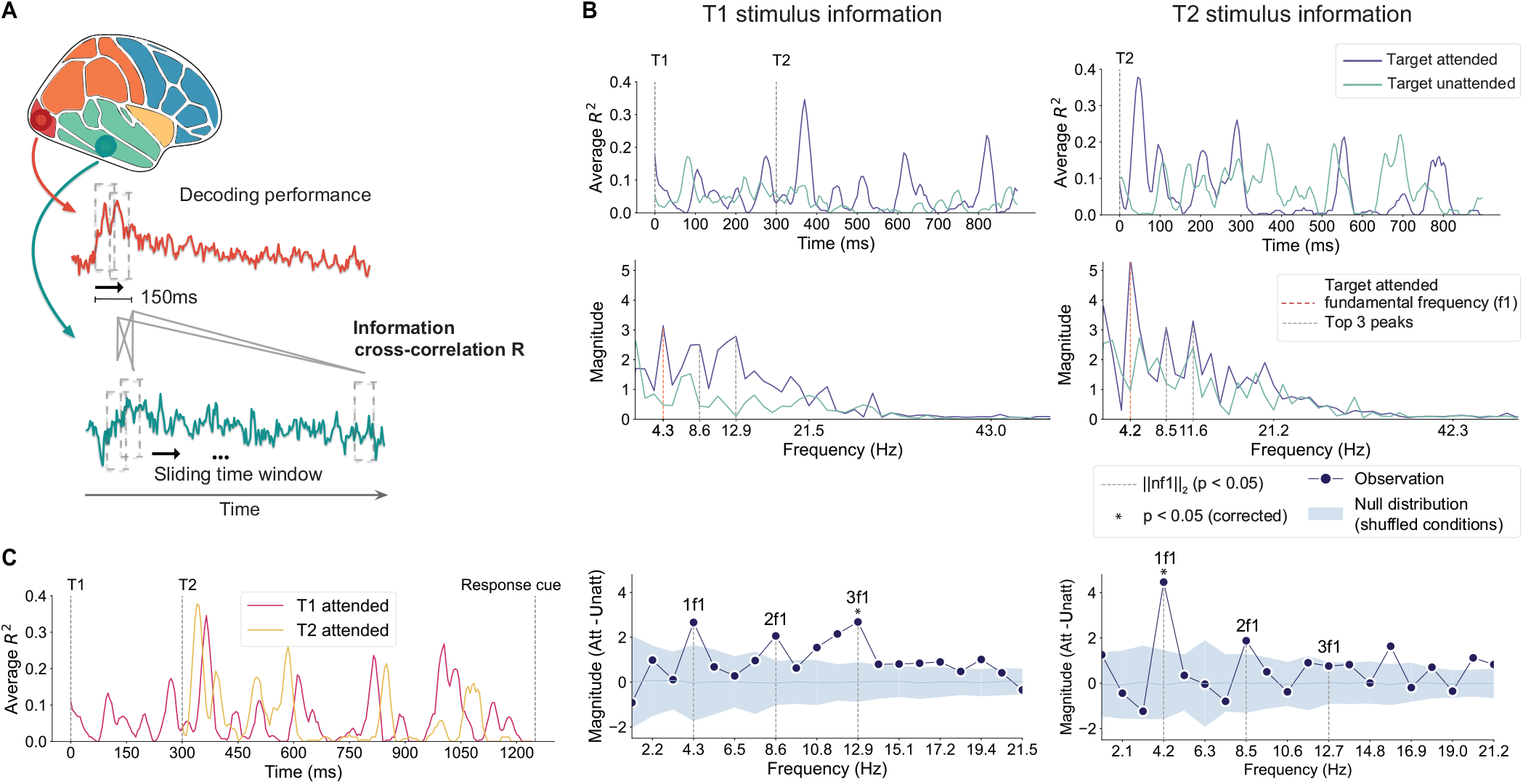
Temporal attention rhythmically modulated sharing of stimulus information between occipital and entorhinal-parahippocampal cortex. **(A)** Schematic of dynamic information cross-correlation R analysis. Stimulus information decoded from occipital cortex (0-150 ms after target onset) are cross correlated with decoded representations in the entorhinal-parahippocampal cortex using sliding windows (5 ms stride and 50 ms window size). **(B)** The *R*_2_ time series (top row) averaged across the 150 ms after target onset for T1 (left) and T2 (right) stimulus information. FFT of the *R*_2_ time series by attention condition (middle row) and their difference (bottom row). The attended *R*_2_ time series exhibited periodicity around a fundamental frequency of 4 Hz for both T1 and T2. **(C)** The *R*_2_ time series for both T1 attended and T2 attended conditions overlaid on the same time axis. It shows that in a hypothetical scenario in which both targets would be attended and there would be no processing trade-offs, the peaks of the *R*_2_ between the occipital and entorhinal–parahippocampal cortices for T1 and T2 roughly coincide.

This observation was confirmed by applying FFTs to the correlation time series and comparing the resulting frequency spectra for attended and unattended conditions. For both targets, the spectra for the attended condition peaked at multiples of 4 Hz, indicating a periodic fluctuation with a 4 Hz fundamental frequency (**Figure 6B, second row**). The total energy of the harmonics (see **Methods**, *Frequency analysis*)—the combined power of the first harmonic (4 Hz) plus the second and third harmonics (8 and 12 Hz)—was significantly greater for attended than for unattended conditions for each target, as established by permutation tests compared to the null distribution (p *<* 0.05).

Overlaying the correlation time series for the T1 attended and T2 attended conditions (**Figure 6C**) shows that the periodic pattern peaks at similar times after T2 onset. This overlap may suggest that, if a participant were attending both targets in a hypothetical scenario with no processing trade-offs, there would be competition between their representations in memory-related regions.

## Discussion

We found that temporal attention to one of two targets (T1 and T2) presented in rapid succession dynamically modulated informational connectivity across cortical networks. These attention-related shifts in connectivity occurred during both early and late time windows and were largely independent of corresponding attentional changes in decoding accuracy. This dissociation indicates that independent of amplifying stimulus representations, temporal attention alters how stimulus information is routed across networks. These findings suggest that temporal attention leads to selective routing, which may serve to resolve competition between stimuli for communication routes. Rather than simply increasing representational strength within a region, temporal attention biases how stimulus information is shared across network pathways at specific moments in time. The attentional modulation of information routing was associated with the processing transition from T1 to T2 and with the theta-rhythmic recurrence of early occipital motifs in memory-related regions.

### Early and late attentional modulations of informational connectivity for each target

Attention modulated informational connectivity network structure mainly at both early and late windows for each target (summarized in **Figure 7**), measured by spatial patterns of closeness centrality. Early modulations occurred ~ 100 ms after each target, around the time when early visual event-related responses (Doherty et al., 2005; Correa et al., 2006) and orientation decoding (Cichy et al., 2015; Wardle et al., 2016; Pantazis et al., 2018) ramp up. Unlike spatial attention (van Es et al., 2018; Dugué et al., 2020; Liu et al., 2021) and feature-based attention (Maunsell and Treue, 2006; Liu et al., 2007; Foster and Ling, 2022), which modulate early visual representations, studies of temporal attention have failed to find early increases in univariate responses or multivariate representations after target onset when temporal expectation is controlled (Zhu et al., 2024). Our results indicate that temporal attention may instead exert early effects without altering local encoding strength by selective routing from visual regions.

**Figure 7.**
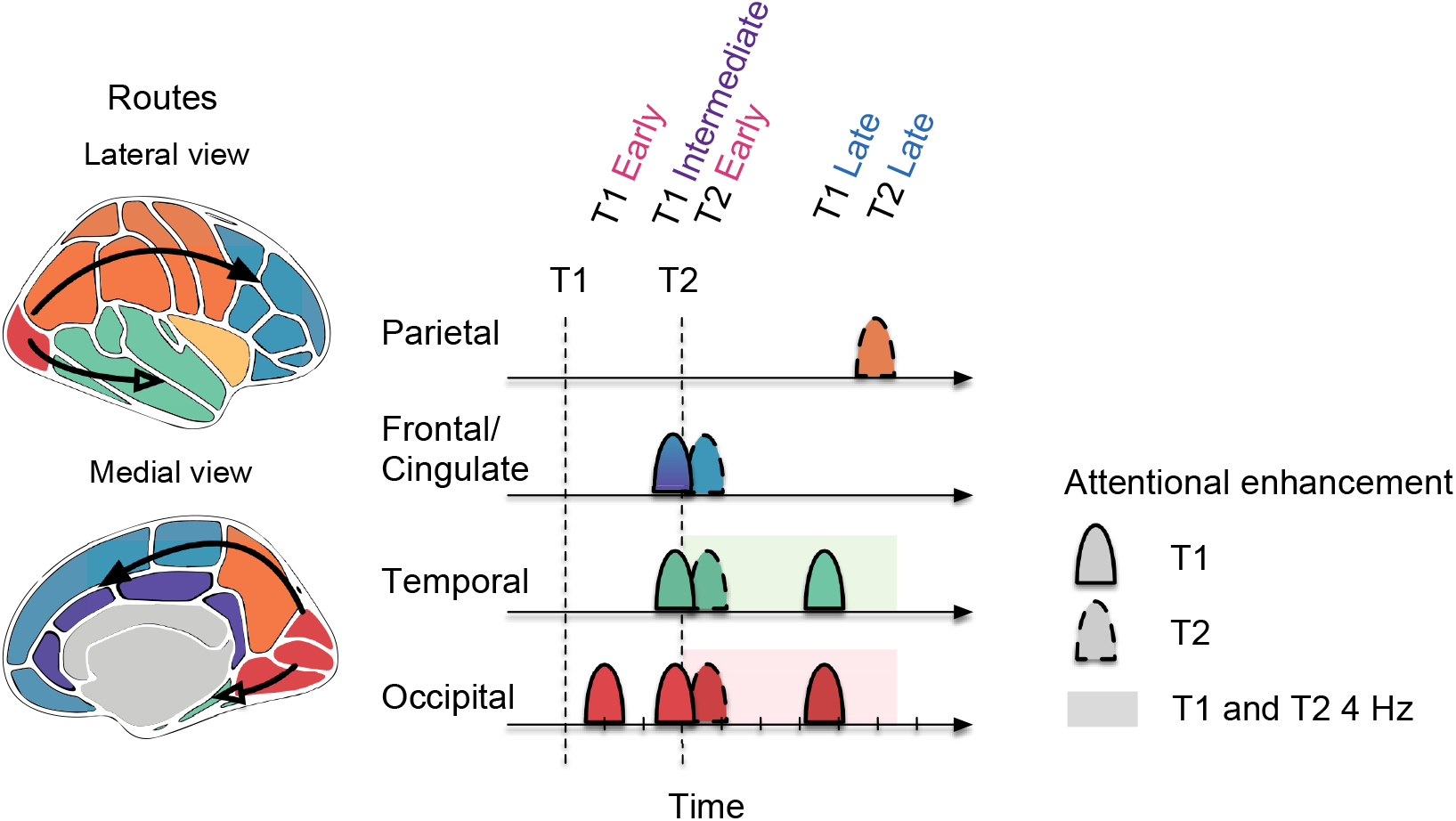
Conceptual illustration of dynamic selective routing. The two proposed pathways are indicated with arrows. In this scheme, competition occurs when T1 and T2 are processed consecutively, and temporal attention modulates the routing of stimulus information from occipital cortex (red) to two downstream areas before decision-making: frontal–cingulate cortex (blue and purple) and temporal cortex (green). The horizontal time axis shows periods when temporal attention enhances informational connectivity (ticks every 100 ms). The midpoint of each marked period of attentional enhancement is aligned with the midpoint of each significant time window. The enhancement periods shown include both significant time windows in which closeness centrality along a route was higher in attended than unattended conditions, as well as windows in which informational connectivity along a route was stronger when a stimulus was attended than when it was unattended (informational connectivity network edges, p *<* 0.05 corrected). For all lobes, T1 and T2 attentional modulations are plotted on the same time axis. The shaded strip along the time axis for occipital and temporal cortex represents the 4 Hz periodic occipital motif in the entorhinal–parahippocampal cortex found for both T1 and T2. Attentional enhancements were left lateralized in all lobes except parietal.

In the early window, T1 showed attentional enhancements of closeness centrality primarily in the occipital lobe (T1 early in **Figure 7**), whereas T2 showed attentional enhancements of closeness centrality in both occipital and temporal lobes (T2 early in **Figure 7**). Thus, even at early latencies traditionally associated with feedforward processing, temporal attention modifies communication structure rather than local gain. Late modulations diverged between targets. In the late time window, T1 showed increased closeness centrality in the occipital and temporal lobes (T1 late in **Figure 7**), but no such effect was significant for T2. Instead, the late window for T2 exhibited a mixture of enhanced closeness centrality in the parietal lobe (T2 late in **Figure 7**) and weak increases in informational connectivity between specific occipital and temporal lobe regions (**Supplementary Figure 4**) that did not survive multiple comparisons correction. One possible account of this variability in late connectivity patterns for T2 is that participants can engage different routing strategies to maintain T2 information until the response window: sharing information with either the temporal lobe (similar to T1) or the parietal lobe. The involvement of the posterior parietal cortex in late T2 routing aligns with its established role in maintaining task-relevant information when visual stimuli are no longer present, in spatial attention-driven and visual working memory tasks (Todd and Marois, 2004; Xu, 2018a; Dugué et al., 2018; Xu, 2024). It has been argued that such maintenance does not only depend on posterior parietal cortex, but selectively uses visual information relayed from occipital-temporal cortex according to task demands (Xu, 2018b). The overall weaker late attentional modulation for T2 may also reflect lower competitive demands during this window, as much of T1 processing may be complete by this time (~ 800 ms after T1). Importantly, this pattern indicates that routing is flexible—attention dynamically allocates information to distinct maintenance systems depending on task timing and competition.

### Pathways for selective routing

The distinct temporal windows of modulation may correspond to distinct pathways that are competitively accessed by both targets. The current findings point to two pathways for selective routing—one during the transition from T1 to T2 involving the fronto-cingulate cortex, and the other after both targets appear involving the temporal lobe (the two pathways are visualized in **Figure 7**).

#### Fronto-cingulate routing during the T1-T2 transition

First, during the transition from T1 to T2, attentional modulations of informational connectivity suggest selective routing to the left fronto-cingulate cortex. In a previous study using the same dataset, we found that temporal attention increased T1 decoding in left fronto-cingulate cortex immediately before T2 onset (Zhu et al., 2024). Here we demonstrate that this time window is also characterized by attention-enhanced informational connectivity associated with T1 (T1 intermediate in **Figure 7**) along left-lateralized pathways involving the fronto-cingulate cortex. Immediately after T2 onset, attended T2 also engaged a left-lateralized pathway extending into the left frontal cortex (T2 early in **Figure 7**), whereas unattended T2 showed delayed and right-lateralized routing. Uncorrected connectivity networks showed that this right-lateralized pathway ultimately crossed over to the left frontal cortex.

We speculate that a possible explanation for these results is that sequential targets compete for access to a left fronto-cingulate bottleneck during the T1 to T2 transition, and temporal attention modulates this routing process. The cingulo-opercular (CO) network has been implicated in selecting a retrocued item and reformatting it into an action-oriented representation during visual working memory tasks (Wallis et al., 2015; Myers et al., 2017). Given that target orientations must be maintained until the response cue in the present task, such a selection and reformatting operation would be relevant here. Thus, temporal attention may resolve competition between T1 and T2 through biased access to large-scale communication pathways.

#### Medial-temporal routing and theta-rhythmic replay

Second, we found evidence for attention-mediated selective routing to memory-related regions in the temporal lobe. Entorhinal cortex and parahippocampal cortex are key regions interconnected with hippocampus within the medial temporal lobe to support memory (Lavenex and Amaral, 2000; Ranganath and Ritchey, 2012; Knierim et al., 2014). Early occipital decoding dynamics appeared to be replayed in these regions rhythmically at 4 Hz throughout the trial. This fundamental frequency falls within the theta band—a frequency range associated with memory-related activity in the medial temporal lobe (Herweg et al., 2020; Lega et al., 2012). Critically, this replay occurred only when stimuli were attended. Both targets showed this periodic replay pattern when attended, with similar absolute timing during the trial (the timing of the periodic replay coincides with T2 early and both T1 and T2 late in **Figure 7**).

These findings suggest that only one target representation can be actively maintained through periodic replay in the medial temporal lobe, and that attention regulates which one is maintained. Thus the medial temporal lobe may be a site of competition for representational resources following the appearance of both targets, leading to selective routing. Together with the frontal–cingulate findings, this observation suggests the presence of at least two bottlenecks in the network: one governing selection during target transition and another governing oscillatory maintenance after stimulus encoding.

### Network communication mechanisms for temporal attentional selection

The early and late attentional modulations observed in the informational connectivity network following target onset suggest that stimulus information is communicated neither instantaneously in a single phase nor continuously over time. Instead, spatial patterns of closeness centrality differed between attended and unattended networks across multiple discrete time windows: for T1 these included early (50–175 ms), intermediate (250–385 ms), and late (625–780 ms) periods; for T2, differences emerged during early (25–215 ms) and late (450–580 ms) windows. This temporal segmentation suggests that stimulus information is transmitted at discrete intervals, each providing a distinct window for network-wide exchange.

One mechanistic interpretation is that stimulus information is conveyed as a “packet,” a term from computer science that refers to a small, formatted unit of data designed to travel efficiently over network connections. Analogous to computer network communication—where a packet can be sent at different times or routed through different paths (Cerf and Kahn, 1974; McQuillan et al., 1980; Gallager, 2003)—a packet of neural information may likewise be transmitted at distinct moments via neural routes. Computer science also provides an interesting, albeit speculative, analogy for selective routing. In the “store-and-forward” mechanism in computer networks (McQuillan and Walden, 1977; Heart et al., 1970), each node buffers and forwards data packets in sequence. Thus information must be buffered in limited-capacity nodes before being relayed downstream.

The present results suggest fronto-cingulate cortex and entorhinal-parahippocampal cortex as two candidate regions that may function as limited-capacity buffers that maintain and forward prioritized stimulus information in the current task. Temporally segmented routing may explain why attentional modulations of informational connectivity appear at multiple, distinct moments rather than as a continuous flow.

Altogether, investigating informational connectivity dynamics and their attentional modulation reveals stimulus processing as a network-level phenomenon. Such a perspective advances understanding of how the brain routes information across large-scale cortical networks during fast-paced, dynamic tasks.

## Conclusion

Here, we provide evidence that temporal attention modulates how, when, and through which pathways stimulus-information travels, rather than simply how strongly it is represented. Beyond enhancing neural responses, temporal attention thus engages a selective routing mechanism to prioritize stimulus information at a specific moment in time. When participants attended to one of two rapidly presented sequential targets, temporal attention modulated informational connectivity across two main pathways: an occipito–frontal–cingulate route involved in selection during the transition between targets, and an occipito–temporal route supporting theta-rhythmic maintenance in the medial temporal lobe. The two sequential targets appeared to compete for access to these routes. Temporal attention modulated informational access to these routes through two complementary mechanisms: discrete bursts of network communication occurring throughout the trial, with a concentration during the transition from the first to the second target, and theta-rhythmic activity that persisted for an extended period following both targets. By identifying routing as the mechanism through which temporal attention resolves sequential competition, this research reveals a network-level account of temporal attention and offers a new framework for understanding how the brain manages information flow during fast, dynamic cognition.

## Methods

### Observers

Ten observers (5 females, mean age = 29 years old, SD = 4 years), including authors RND and KJT, participated in the study. Each observer first completed one behavioral training session, followed by two separate 2-hour MEG recording sessions conducted on different days, yielding a total of 20 MEG sessions. Collecting two sessions per observer enabled us to assess within-observer reliability across sessions and follows an approach used in other MEG studies of vision to collect multiple sessions per observer (Kok et al., 2017; Besserve et al., 2007). Experimental protocols were approved by the University Committee on Activities involving Human Subjects at New York University. All observers had normal or corrected-to-normal vision using MR safe lenses. All observers provided informed consent.

### Stimuli

Stimulus presentation was programmed in MATLAB using Psychtoolbox (Brainard and Vision, 1997; Pelli and Vision, 1997; Kleiner et al., 2007) and run on an iMac. During MEG acquisition, images were delivered with an InFocus LP850 projector (Texas Instruments, Warren, NJ) and reflected by a mirror onto a translucent screen (1024 × 768 pixels; 60 Hz) positioned 42 cm from the participant. Stimuli appeared on a mid-gray background (206 cd/m^2^), and timing accuracy was verified with a photodiode. For the behavioral training conducted outside the MEG, stimuli were shown on a gamma-corrected Sony Trinitron G520 CRT monitor (1024 × 768 pixels; 60 Hz) at a viewing distance of 56 cm. Participants maintained a stable head position using a combined chin-and-head rest.

#### Visual targets

Targets were foveally presented, full-contrast sinusoidal gratings (1.5 cycles/deg). Each grating subtended 4° in diameter and was windowed with a raised-cosine smooth taper so that the outermost 0.4° gradually fell to zero contrast.

#### Auditory cues

Both precues and response cues were 100-ms pure-tone sinusoids, gated with 10-ms cosine ramps at onset and offset to minimize audible transients. A high tone (1046.5 Hz; C6) signaled T1, whereas a low tone (440 Hz; A4) signaled T2.

### Task

Observers were instructed to voluntarily direct their temporal attention to a specific time point and report the tilt of the cued target. On each trial (ref to **Figure 1A**), two visual targets (T1 and T2) were presented sequentially at the same spatial location. Each target appeared for 50 ms, with a 300 ms stimulus onset asynchrony (SOA) between them—a timescale known to elicit temporal attentional tradeoffs in prior psychophysical work (Denison et al., 2017, 2021; Fernández et al., 2019; Palmieri and Carrasco, 2024). Each target was slightly tilted either clockwise (CW) or counterclockwise (CCW) relative to either the vertical or horizontal axis (**Figure 1B**); tilt direction and axis were manipulated independently and counterbalanced for each target.

Each trial began with an auditory precue occurring 1,050 ms prior to target presentation that instructed observers to allocate attention to T1 or T2 (high vs. low tone). A second auditory cue, presented 950 ms after the targets, specified which target should be reported. Participants indicated the tilt direction (CW vs. CCW) using a two-button response within 1,500 ms. Feedback was conveyed at the end of the trial by changing the color of the fixation marker (green for correct, red for incorrect, blue for timeout).

On each trial, the timing of target presentation was fully predictable following the auditory precue. The attended target varied from trial to trial based on the precue, while the target designated for behavioral report was determined by a separate response cue presented at the end of each trial. In 80% of trials, the precue directed attention to a single target (T1 or T2), and in those cases, the response cue matched the precued target 75% of the time—creating an incentive for participants to follow the precue. Both the identity of the precued target and the cue validity (match vs. mismatch of precue and response cue) were randomized across trials. Each MEG session included 192 trials each for precue T1 and precue T2 conditions. Trials were labeled as “attended” with respect to the precued target and “unattended” with respect to the non-target on that trial, resulting in 192 attended and 192 unattended trials per target. Note that the response cue allowed us to verify that behavior depended on cue validity; however, because the response cue was presented after both targets at the end of the trial, it was irrelevant to the conditions used for decoding. Additionally, the identity of the auditory precue (high vs. low tone) was orthogonal to the stimulus orientation (vertical or horizontal) and thus carried no information relevant to decoding the visual representation.

The experiment also included neutral trials (20% of all trials), which were cued with a combination of the high and low tones, providing no temporal information and thus indicating that attention should be distributed across both targets. These trials enabled behavioral assessment of temporal attentional selectivity (see **Supplementary Figure 1**). Due to their reduced count—50% fewer trials relative to the precue-T1 and precue-T2 conditions—neutral trials were excluded from MEG decoding analyses to avoid trial-count imbalances that could bias classification performance.

#### Training

Prior to MEG recording, participants completed a behavioral training session to familiarize them with the task and calibrate individual tilt discrimination thresholds. Tilt magnitude was adjusted separately for each observer using a 3-up-1-down staircase procedure to target approximately 79% accuracy on neutral trials (mean threshold tilt = 0.76° across observers).

### Eye tracking

Throughout each trial, observers fixated on a central circular marker (0.15° diameter). Eye position was continuously recorded at 1000 Hz using an EyeLink 1000 system (SR Research). Calibration was performed at the beginning of each session using a five-point grid, which allowed gaze coordinates to be mapped to degrees of visual angle.

### MEG acquisition

Each MEG session consisted of 12 experimental blocks, each lasting approximately 6 minutes. Observers were allowed to rest between blocks and resumed the task at their own pace by pressing a button to continue.

Prior to MEG acquisition, individual head shapes were digitized using a FastSCAN laser scanner (Polhemus, VT, USA). Fiducial points were marked digitally at the nasion, forehead, and bilaterally at the tragus and peri-auricular locations. The positions of these reference points were recorded at both the beginning and end of each session. To enable co-registration of head position with MEG sensor locations, electrodes were affixed at the forehead and peri-auricular sites corresponding to the digitized fiducials.

MEG signals were acquired continuously using a 157-channel axial gradiometer system (Kanazawa Institute of Technology, Kanazawa, Japan) housed in the KIT/NYU MEG facility at New York University. Background magnetic interference was tracked using three orthogonal reference magnetometers placed approximately 20 cm from the sensor array. Data were sampled at 1000 Hz, with online DC correction and a high-pass filter set at 200 Hz.

### Preprocessing

MEG preprocessing was conducted in MATLAB using the FieldTrip toolbox for EEG/MEG analysis (Oostenveld et al., 2011), following these steps: 1) All trials were visually examined, and those containing eye blinks or other artifacts were manually excluded. Across sessions, the number of rejected trials ranged from 18 to 88 (3.49–17.05%), with a mean of 51.75 trials (10.03%) and a standard deviation of 20.06. 2) Channels exhibiting abnormally high or low variability were automatically flagged based on their time series standard deviations. 3) Time series from these noisy channels were replaced by spatial interpolation from neighboring sensors. The number of interpolated channels per session ranged from 0 to 6 (0–3.82%), with a mean of 3.85 (2.45%) and a standard deviation of 1.50. 4) To remove environmental noise, signals from the reference magnetometers were linearly regressed out of the MEG sensor time series.

### Source reconstruction

We performed source reconstruction using MNE Python (Gramfort et al., 2013). For each participant, we generated a cortical surface mesh from their anatomical MRI with 4,000 vertices per hemisphere. Coregistration between MEG and MRI data was performed automatically using three fiducial landmarks (nasion and bilateral peri-auricular points) and digitized scalp points acquired via laser scanning (Gramfort et al., 2013; Houck and Claus, 2020). Forward models were calculated using a single-shell Boundary Element Model (BEM) to approximate the head’s geometry and tissue conductivities. Source estimates were computed by inverting the forward model using dynamic statistical parametric mapping (dSPM) (Dale et al., 2000). At each vertex, the estimated source activity was modeled as a dipole oriented perpendicular to the cortical surface. The sign of the dipole reflects current direction—positive for outward, negative for inward flow (Wang et al., 2023). In dSPM-based source localization, the typical Dipole Localization Error (DLE) is approximately 2 cm, and the Spatial Dispersion (SD) is around 4 cm (Hauk et al., 2011; Hedrich et al., 2017). DLE represents the Euclidean distance between the true dipole location and the maximum of the estimated source map, while SD captures the extent to which the estimated activity is spatially spread around the true source location (Molins et al., 2008; Hauk et al., 2011).

Cortical parcellation was performed using the 34 regions per hemisphere defined in the Desikan–Killiany (DK) atlas (Desikan et al., 2006). These regions of interest (ROIs) can be grouped into broader anatomical lobes (occipital, parietal, temporal, frontal, cingulate and insula) following the mapping scheme in (Klein and Tourville, 2012).

### Decoding

To decode stimulus orientation (vertical vs. horizontal), we trained linear support vector machine (SVM) classifiers independently at each time point (Cichy et al., 2015; King and Dehaene, 2014). A 5-fold cross-validation was used to partition trials into training and testing sets, ensuring unbiased estimates of classification performance. Decoding was performed separately for each target (T1 and T2) under each precue condition, resulting in time-resolved decoding accuracy time series. For instance, in decoding T1 orientation, trials with a precue to T1 were labeled as attended, while those cued to T2 were considered unattended. To improve signal-to-noise ratio, we generated pseudotrials by averaging across small groups of 5 trials (Isik et al., 2014; Meyers, 2013; Wardle et al., 2016), and also averaged the data over 5 ms temporal windows (Isik et al., 2014). The entire decoding process was repeated 100 times using randomized pseudotrial assignments to mitigate any variability introduced by the averaging procedure.

In source space, we performed decoding of stimulus orientation using estimated source activations within anatomically defined ROIs from the DK atlas (see *Source reconstruction*). Each ROI contained multiple vertices, and the corresponding time series were derived from the source reconstruction output. Because the number of features (vertices) within an ROI often exceeded the number of available samples (trials), we applied dimensionality reduction to prevent overfitting. Specifically, for ROIs containing more than 100 vertices, we employed univariate feature selection using an ANOVA F-test (Pedregosa et al., 2011; Gramfort et al., 2013), applied only to the training set within the 5-fold cross-validation. The top 100 vertices with the highest F-scores were selected as input features for the classifier, ensuring that the feature dimensionality did not exceed 100 for any ROI.

To obtain decoding performance for larger regions (e.g., occipital and entorhinal–parahippocampal cortex), we averaged decoding accuracy across the constituent ROIs within each region.

### Dynamic informational connectivity analysis

#### Network construction

To obtain higher signal-to-noise stimulus information, we averaged decoding accuracy time series from source-reconstructed atlas-based regions (*N* = 68 regions, or 34 per hemisphere) across all sessions per attention condition to form a “super-subject” and conducted all further analyses using these data. We estimated informational connectivity networks based on the decoding accuracy timeseries of the “super-subject”. We performed this analysis across sliding time windows to understand how informational connectivity changes across time, a dynamic informational connectivity analysis. Formally, each region *i* ∈ *V* = {1, …, *N*} was treated as a node in a graph. For each time *t*, we estimated a weighted undirected graph *G*(*t*) = (*V, E*, **Z**(*t*)), where *E* = (*i, j*) is the set of all possible undirected edges between region *i* and another region *j* ∈ *V*, **Z**(*t*) = [*Z*_*ij*_(*t*)] is the adjacency matrix encoding informational connectivity.

For each time *t*, we define a sliding window of width Δ*t*, denoted *W* (*t*) = [*t, t* +Δ*t*], where Δ*t* contains discrete time steps *τ*_*l*_ ∈ {*τ*_1_, …, *τ*_*m*_}. Here, each time step *τ*_*l*_ was defined as a 5 () ms bin (see details in *Decoding*) and the window duration was 50 ms (i.e., *m* = 10). Within each window *W* (*t*), the stimulus orientation decoding accuracies for each region *i* are denoted by **a**_*i*_(*t*) = *a*_*i*_(*t* + *τ*_1_), *a*_*i*_(*t* + *τ*_2_), …, *a*_*i*_(*t* + *τ*_*m*_) ∈ ℝ ^*m*^.

Informational connectivity between regions *i* and *j* at time *t* is defined as the Pearson correlation between their accuracy vectors using **Equation 1**:

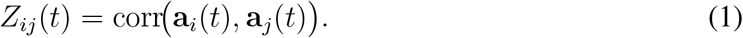

Thus, by sliding the window across time, we generate a time-varying network {*G*(*t*)}, which captures the evolution of informational connectivity across conditions.

#### Closeness Centrality

Closeness centrality is a graph-theoric measure that summarizes the distance between one node in the network and all other nodes and quantifies how efficiently signals can spread through a network (Freeman et al., 1979). Here we used correlation distances to calculate closeness centrality for each atlas-based region, providing a summary of each region’s shared information with all other regions in each sliding time window, and thus the centrality of that region within the informational connectivity network across time.

For each region *i* ∈ *V*, we define its closeness centrality with respect to the informational connectivity network *G*(*t*) in **Equation 2**:

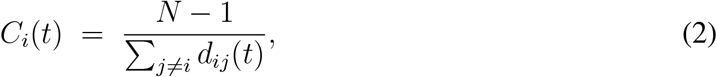

where *d*_*ij*_(*t*) denotes the shortest correlation distance between regions *i* and *j*, given by **Equation 3**:

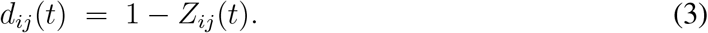

Thus, *C*_*i*_(*t*) represents the average inverse correlation distance from region *i* to all other regions, with larger values indicating stronger and more efficient information flow.

For a given lobe, the closeness centrality was calculated as the L2-norm of the centrality values for each ROIs within the lobe, providing a measure of overall centrality strength.

#### KL Divergence

Kullback-Leibler (KL) divergence is an information-theoretic measure that summarizes how different one probability distribution is from another. In statistics and machine learning, it is used to quantify the information loss incurred when an approximate distribution is used in place of a reference distribution (Kullback and Leibler, 1951; Cover, 1999). Here, we use the distribution of closeness centrality to summarize global network structure. We then use KL divergence to compare network structure across attention conditions within each sliding time window, providing a summary of how strongly the attended and unattended informational connectivity networks diverge over time.

Let 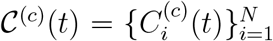 denote the set of region-wise closeness centrality values at time *t* under condition *c*. We normalize these values to obtain a distribution of scaled closeness centrality values across regions using **Equation 4**:

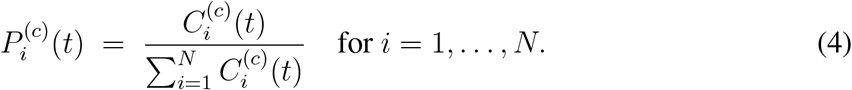

To compare the attended (*c* = att) and unattended (*c* = unatt) networks, we compute the KL divergence, which is a relative entropy measure of the difference between two distributions using **Equation 5**:

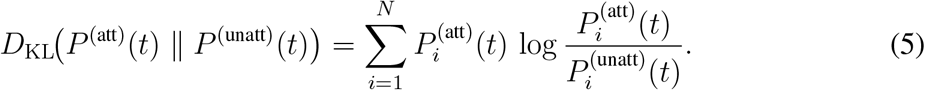

### Information cross-correlation (time-lagged connectivity)

We used a cross-correlation approach to investigate whether an early motif in one region recurred in other regions.

For region *i*, consider sliding windows *W* (*t*_*i*_) with *t*_*i*_ ∈ 𝒯_*i*_ = [0, 150] ms after target onset; for region *j*, windows *W* (*t*_*j*_) with *t*_*j*_ ∈ 𝒯_*j*_. Let **a**_*i*_ *W* (*t*_*i*_) and **a**_*j*_ *W* (*t*_*j*_) denote the vectors of decoding accuracies extracted within those windows. Consistent with the sliding window used in previous sections, the window duration was 50 ms.

The cross-correlation between window *W* (*t*_*i*_) and *W* (*t*_*j*_) is using **Equation 6**:

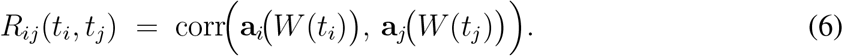

We use Pearson correlation and it can expand explicitly as in **Equation 7**:

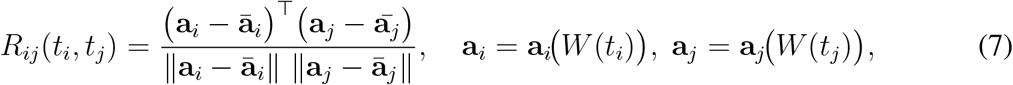

with **ā**_*i*_ and **ā**_*j*_ the within-window means. The collection {*R*_*ij*_(*t*_*i*_, *t*_*j*_)} forms an |𝒯_*i*_| × |𝒯_*j*_| cross-correlation matrix for the pair (*i, j*).

Here, for each sliding time window within the first 150 ms after target onset in the occipital cortex (region *i*), the cross-correlation with each sliding time window in the entorhinal–parahippocampal cortex (region *j*) is denoted by *R* (*R* = *R*_*ij*_(*t*_*i*_, *t*_*j*_)). To summarize the early occipital pattern, we averaged *R*_*ij*_(*t*_*i*_, *t*_*j*_)^2^ across *t*_*i*_ for each fixed *t*_*j*_, thereby collapsing the |T_*i*_| × |𝒯_*j*_| matrix into a length-|𝒯_*j*_| time series of averaged *R*^2^ values for region *j*. Formally, we can use **Equation 8**:

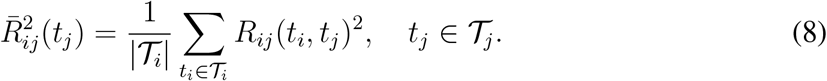

#### Frequency analysis

For both T1 and T2, the averaged *R*^2^ values for the attended condition within the first 150 ms after target onset in the occipital cortex has some periodic patterns in the entorhinal–parahippocampal cortex over time. We then applied FFT analysis for the *R*^2^ time series to quantify such periodicity.

We characterized periodic patterns in cross-correlation time series between regions of interest using a frequency analysis. Whereas a pure sinusoidal periodic waveform is characterized by a single frequency, a non-sinusoidal periodic waveform also has harmonics that reflect the strength of the periodic signal. The lowest frequency component of a periodic signal is the fundamental frequency (*f*_0_), which determines the signal’s basic repetition rate. Harmonics refer to sinusoidal components of a signal that occur at integer multiples of a fundamental frequency (*f*_0_) as shown in **Equation 9**:

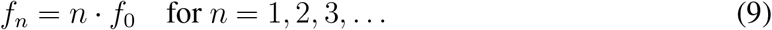

*f*_1_ = *f*_0_ (first harmonic), *f*_2_ = 2*f*_0_ (second harmonic), *f*_3_ = 3*f*_0_ (third harmonic), etc.

Here, we found a 4 Hz fundamental frequency, with 8 Hz and 12 Hz as its second and third harmonic frequencies, respectively, for both the T1 and T2 attended conditions. Specifically, we found 3 prominent peaks in the FFT and let the lowest frequency (*f*_1_) be the fundamental frequency. The other frequencies of the peaks are the multiplied frequencies for the lowest frequency (i.e., 2*f*_1_, 3*f*_1_).

To quantify the overall strength of the periodic signal, we calculated harmonic energy as the L2-norm of harmonic amplitudesv 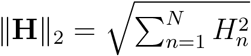, where *H*_*n*_ = |*X*(*nf*_1_)| is the magnitude of the Fourier transform *X*(*f*) of the signal at the *n*-th harmonic frequency.

Here we simplified the notation by letting the amplitude of the *n*-th harmonic to be *n*f1, and the harmonic energy to be 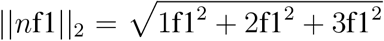 with *n* = 3 (Appelbaum et al., 2010).

### Statistical analysis

#### General procedure

##### Null distribution construction

To assess the statistical significance of the difference between attention conditions, we used nonparametric methods for the informational connectivity network. For a given measure (e.g., closeness centrality) we generated null distributions for the difference between attention conditions by randomly swapping or not swapping the condition labels (attended or unattended) session by session (*n* = 20 sessions) for the decoding accuracy timeseries. We call these conditions with shuffled labels “pseudo-conditions.” To generate a null distribution for the effect of attention on a given measure of interest, we calculated the measure for each pseudo-condition in the same way as we had for the original conditions with correct labels, took the difference between “attended” and “unattended” pseudo-conditions, and repeated this randomization and analysis procedure 1000 times. For each target, the time window of interest was 0–900 ms after the target onset, so the time range was not extended to the response cue and was comparable across targets

##### Non-parametric cluster correction

We used a non-parametric cluster correction for the time series of a given measure of interest, with measure-specific cluster correction methods described below. Significant time windows were identified as successive sliding windows in which informational connectivity measures survived cluster corrections. To do this, we first construct a null distribution of cluster statistics. The cluster-level statistic for each permutation was calculated as follows:

1. For sample at each time step, get the difference of a given measure of interest (e.g., closeness centrality) between two pseudo-conditions as the permuted value. The test statistic is obtained by comparing the permuted value with the null distribution, previously generated for each time step as described in *Null distribution construction* or constructed more conservatively on empirical informational connectivity network. For more information on the null distribution model in network see Váša and Mišić (2022).
2. Select all samples whose test statistic is larger than some threshold. Higher thresholds are better suited for identifying stronger, short-duration effects, whereas lower thresholds are better suited for identifying weaker, long-duration effects (Maris and Oostenveld, 2007). For example, threshold= 0.85 indicates that the permuted value is greater than the 85th percentile of the null distribution. Measure-specific thresholds are described in their respective sections below.
3. Cluster the selected samples into connected sets on the basis of temporal adjacency.
4. Calculate cluster-level statistics by taking the sum of the test statistic within a cluster.
5. Take the largest of the cluster-level statistics.

##### Reporting the significant windows

In reporting significant time windows in the text, we give the starting time point of the first sliding window and the end time point of the last sliding window within the cluster. Visualizations shade the interval between the starting time points of the first and the last sliding windows to better match the plotted timeseries, where the value plotted at each time point represents the value derived from the 50 ms sliding window starting at that time point.

#### Closeness centrality

To assess the effect of temporal attention on closeness centrality across the full time series and identify time windows with significant differences between attention conditions, we used a non-parametric test with cluster correction as described in *Non-parametric cluster correction*. For the cluster-level statistic in each permutation, the null distribution generated for each time step is described in the *Null distribution construction* The reported significant clusters are corrected with threshold values of 0.85, 0.95, or 0.99 (see figure captions).

#### Informational connectivity network edges

To assess the statistical significance of observed differences in informational connectivity network edges between attention conditions, we computed test statistics by comparing the observed values with the null distribution for each edge. Multiple comparisons across edges were then controlled using the Benjamini–Hochberg false discovery rate procedure. The edges for a specific time window are constructed based on the averaged decoding accuracies across the time window.

#### Cross-correlation frequency energy

We sought to determine whether the cross-correlation time-series exhibited greater periodicity when a stimulus was attended vs. unattended. To do so, we assessed whether the harmonic energy of the major spectral peaks (4 Hz, 8 Hz, 12 Hz) differed between attention conditions, beyond what would be expected from a null distribution. We subtracted the unattended FFT spectrum from the attended ones and calculated the energy of the harmonics of this FFT spectrum difference. We then compared the observed energy of the harmonics to a null distribution as described in *General procedure*.

#### KL divergence

We investigated how fluctuations in KL divergence from attended to unattended conditions across time deviated from the time-series of average KL divergence. To correct the test statistics for the full time series of the KL divergence, we used the same nonparametric cluster correction as described above *Non-parametric cluster correction*, using a conservatively constructed null distribution centered on the averaged KL value. The null distribution of networks is more conservative when constructing the null model from the empirical network (Váša and Mišić, 2022). Specifically, for each attention condition, we shuffled the empirical distribution of closeness centrality across time to construct a pseudo-condition. We then obtained the KL divergence between the two pseudo-conditions. The null distribution of the KL divergence was generated by repeating this randomization and analysis procedure 1000 times. Therefore, for each time step, we have a histogram of 1000 KL values as a null distribution. Significant clusters reported are corrected with a fixed threshold of 0.6 for both T1 and T2. The significant cluster showed the time periods where the KL divergence is greater than the time-series of average KL divergence.

## Acknowledgments

We thank Sirui Liu and Luis Ramirez for assistance with early versions of the experiment. We thank Hsin-Hung Li as well as members of the Denison and Carrasco Labs for helpful comments. We thank Jeffrey Walker at the NYU MEG lab for technical assistance. This research was supported by National Institutes of Health National Eye Institute R01 EY037358 and F32 EY025533 to R.N.D., R01 EY019693 to M.C., T32 EY007136 to NYU, National Defense Science and Engineering Graduate Fellowship to K.J.T., and by funding from Boston University to R.N.D.

## Supplementary Figures and Tables

**Supplementary Table 1.** Node number for 68 atlas-based ROIs.

| ROI# | Occipital | ROI# | Parietal | ROI# | Temporal |
| --- | --- | --- | --- | --- | --- |
| 1 | left lateral occipital | 9 | left superior parietal | 19 | left superior temporal |
| 2 | right lateral occipital | 10 | right superior parietal | 20 | right superior temporal |
| 3 | left lingual | 11 | left inferior parietal | 21 | left middle temporal |
| 4 | right lingual | 12 | right inferior parietal | 22 | right middle temporal |
| 5 | left cuneus | 13 | left supramarginal | 23 | left inferior temporal |
| 6 | right cuneus | 14 | right supramarginal | 24 | right inferior temporal |
| 7 | left pericalcarine | 15 | left postcentral | 25 | left bankssts |
| 8 | right pericalcarine | 16 | right postcentral | 26 | right bankssts |
|  |  | 17 | left precuneus | 27 | left fusiform |
|  |  | 18 | right precuneus | 28 | right fusiform |
| ROI# | Temporal | ROI# | Frontal | ROI# | Frontal |
| 29 | left transverse temporal | 37 | left superior frontal | 45 | left pars opercularis |
| 30 | right transverse temporal | 38 | right superior frontal | 46 | right pars opercularis |
| 31 | left entorhinal | 39 | left rostral middle frontal | 47 | left pars orbitalis |
| 32 | right entorhinal | 40 | right rostral middle frontal | 48 | right pars orbitalis |
| 33 | left temporal pole | 41 | left caudal middle frontal | 49 | left lateral orbitofrontal |
| 34 | right temporal pole | 42 | right caudal middle frontal | 50 | right lateral orbitofrontal |
| 35 | left parahippocampal | 43 | left pars triangularis | 51 | left medial orbitofrontal |
| 36 | right parahippocampal | 44 | right pars triangularis | 52 | right medial orbitofrontal |
| ROI# | Frontal | ROI# | Cingulate | ROI# | Insula |
| 53 | left precentral | 59 | left isthmus cingulate | 67 | left insula |
| 54 | right precentral | 60 | right isthmus cingulate | 68 | right insula |
| 55 | left paracentral | 61 | left posterior cingulate |  |  |
| 56 | right paracentral | 62 | right posterior cingulate |  |  |
| 57 | left frontal pole | 63 | left rostral anterior cingulate |  |  |
| 58 | right frontal pole | 64 | right rostral anterior cingulate |  |  |
|  |  | 65 | left caudal anterior cingulate |  |  |
|  |  | 66 | right caudal anterior cingulate |  |  |

**Supplementary Figure 1.**
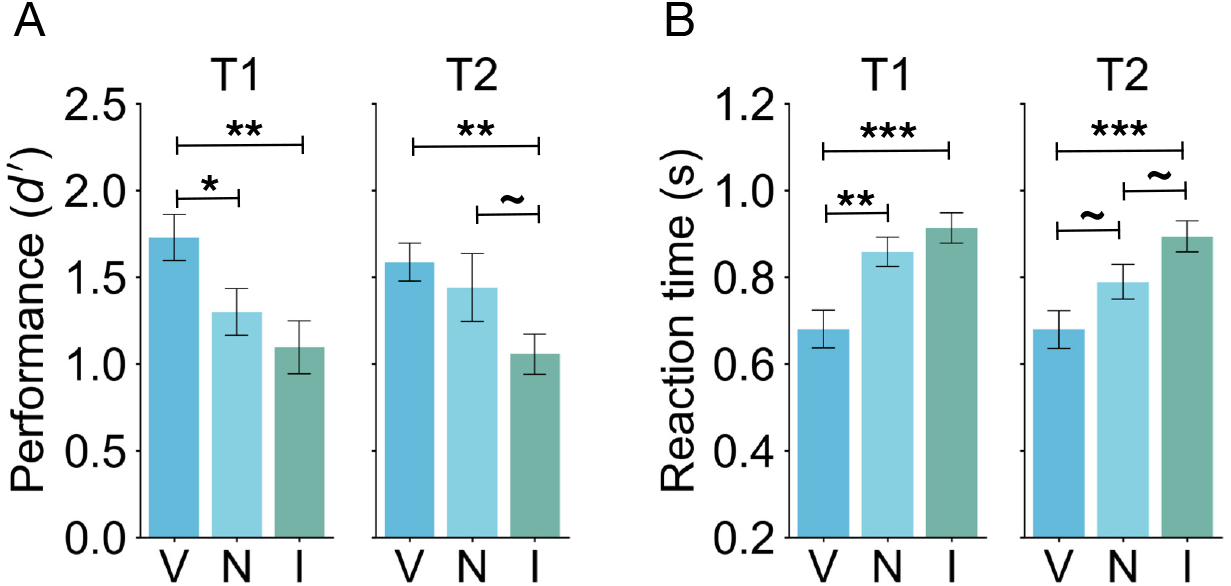
Behavioral results including the neutral condition. Perceptual sensitivity was impaired in the neutral condition (when the temporal precue was uninformative), relative to the valid condition for T1. This result confirms that observers could not fully process both stimuli and instead relied on attention to select the more relevant stimulus in the sequence. Such behavioral benefits are consistent with previous studies using the two-target temporal cueing task (Denison et al., 2017, 2021; Fernández et al., 2019; Duyar et al., 2024), indicating that this task reliably elicits temporal attentional selection. **(A)** Tilt discrimination (sensitivity) and **(B)** reaction time for each target (T1, T2) and validity condition (V = valid, N = neutral, I = invalid). Error bars indicate ±1 SEM. ~ p *<* 0.1, * p *<* 0.05, ** p *<* 0.01; *** p *<* 0.001.

**Supplementary Figure 2.**
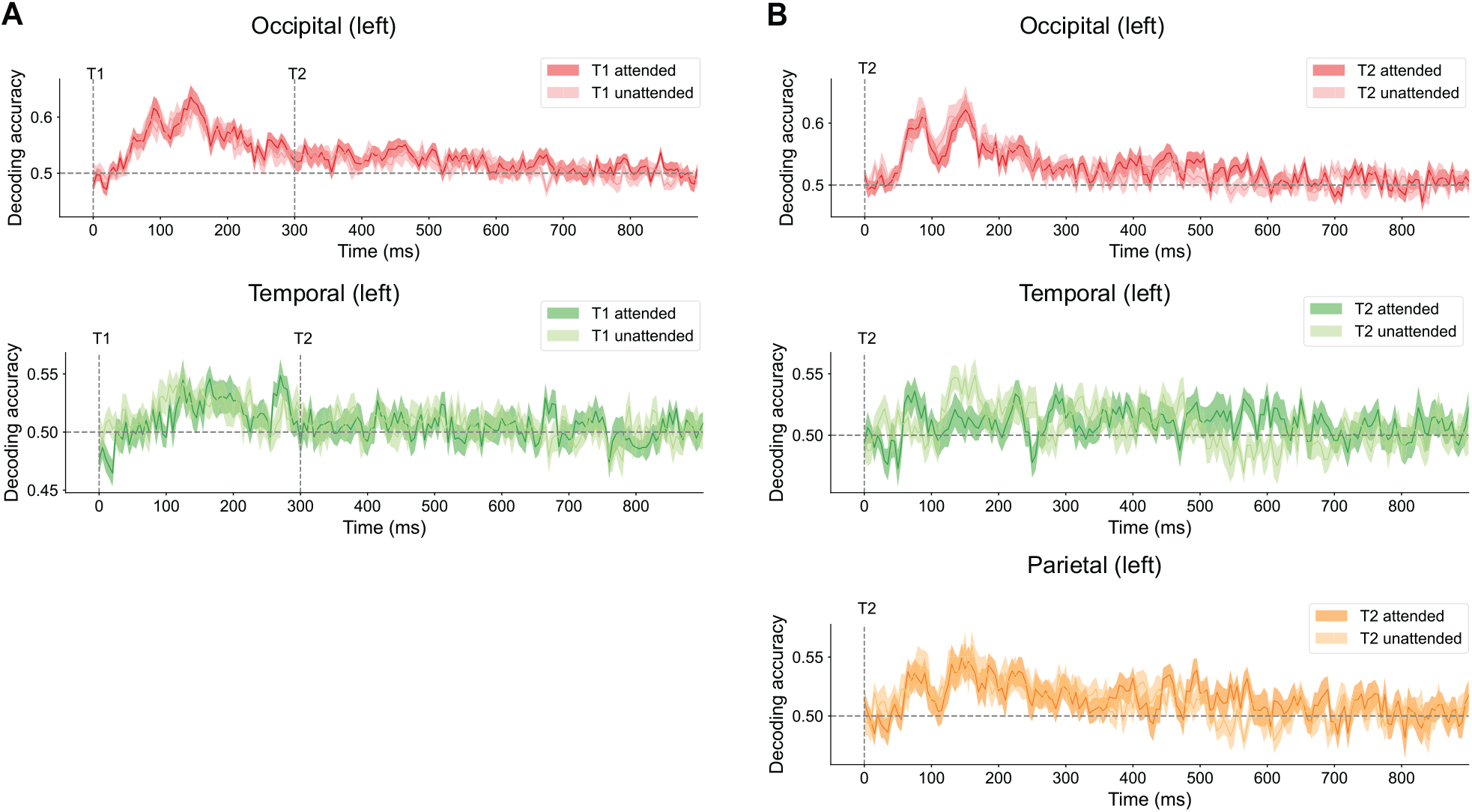
Decoding accuracy for target attended vs target unattended in different lobes. **(A)** Decoding accuracies for T1 in lobes that show enhancement of closeness centrality in **Figure 3. (B)** Decoding accuracies for T2 in lobes that show enhancement of closeness centrality in **Figure 4**.

**Supplementary Figure 3.**
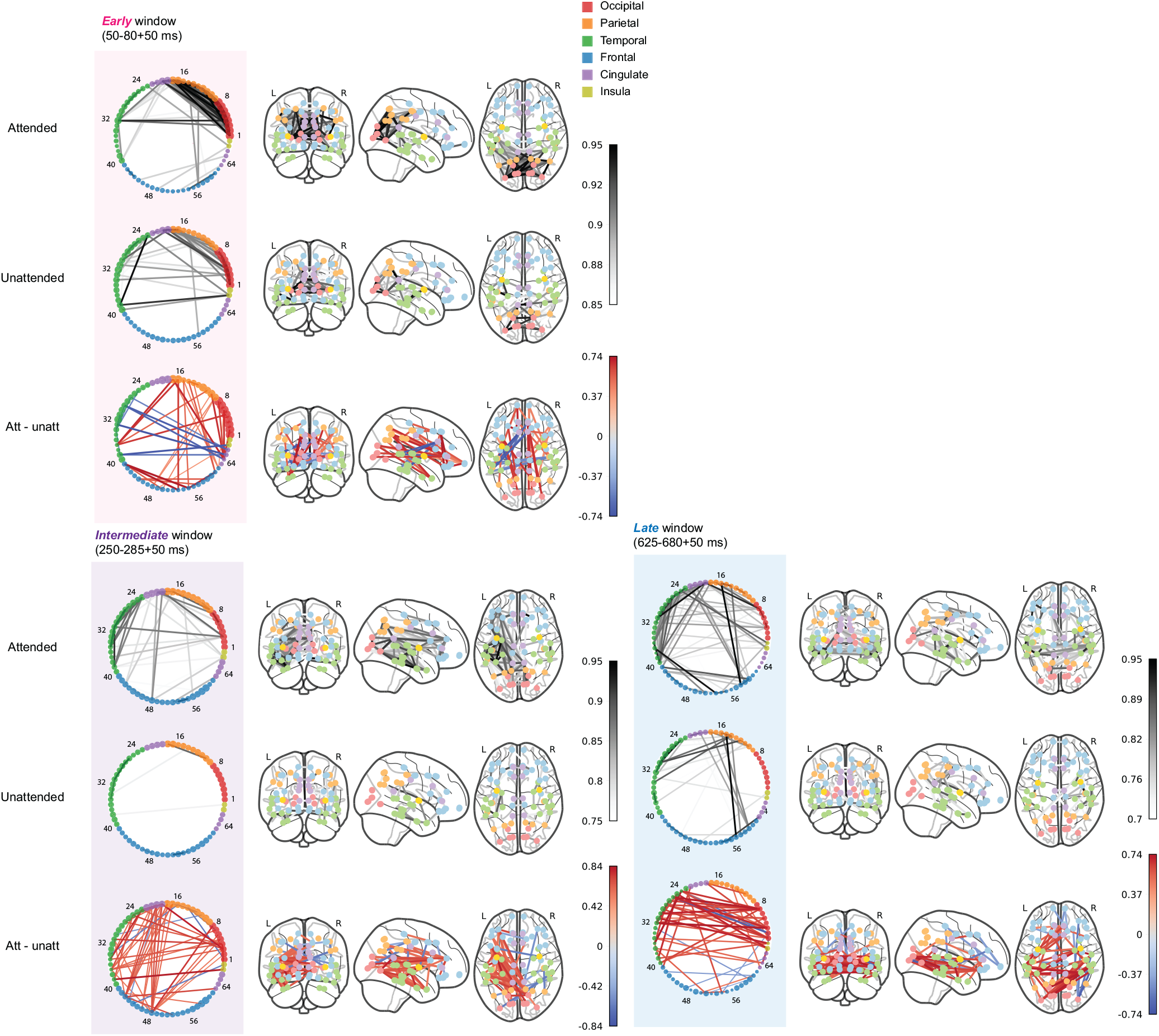
Visualization for averaged dynamic informational connectivity networks for T1 at different critical time windows. For each time window, the network is shown as a chord diagram in which each node represents an ROI; corresponding coronal, sagittal, and axial views are shown from left to right. The visualization of target attended networks is on the first row, and target unattended networks are on the second row. The darker the edges between the nodes, the stronger the connectivity is between them. The third row shows the difference in informational connectivity between attended and unattended conditions. The presented edges are p *<* 0.01 compared to a shuffled null distribution. Edges with higher values are shown in red, indicating greater connectivity during the attended than the unattended condition.

**Supplementary Figure 4.**
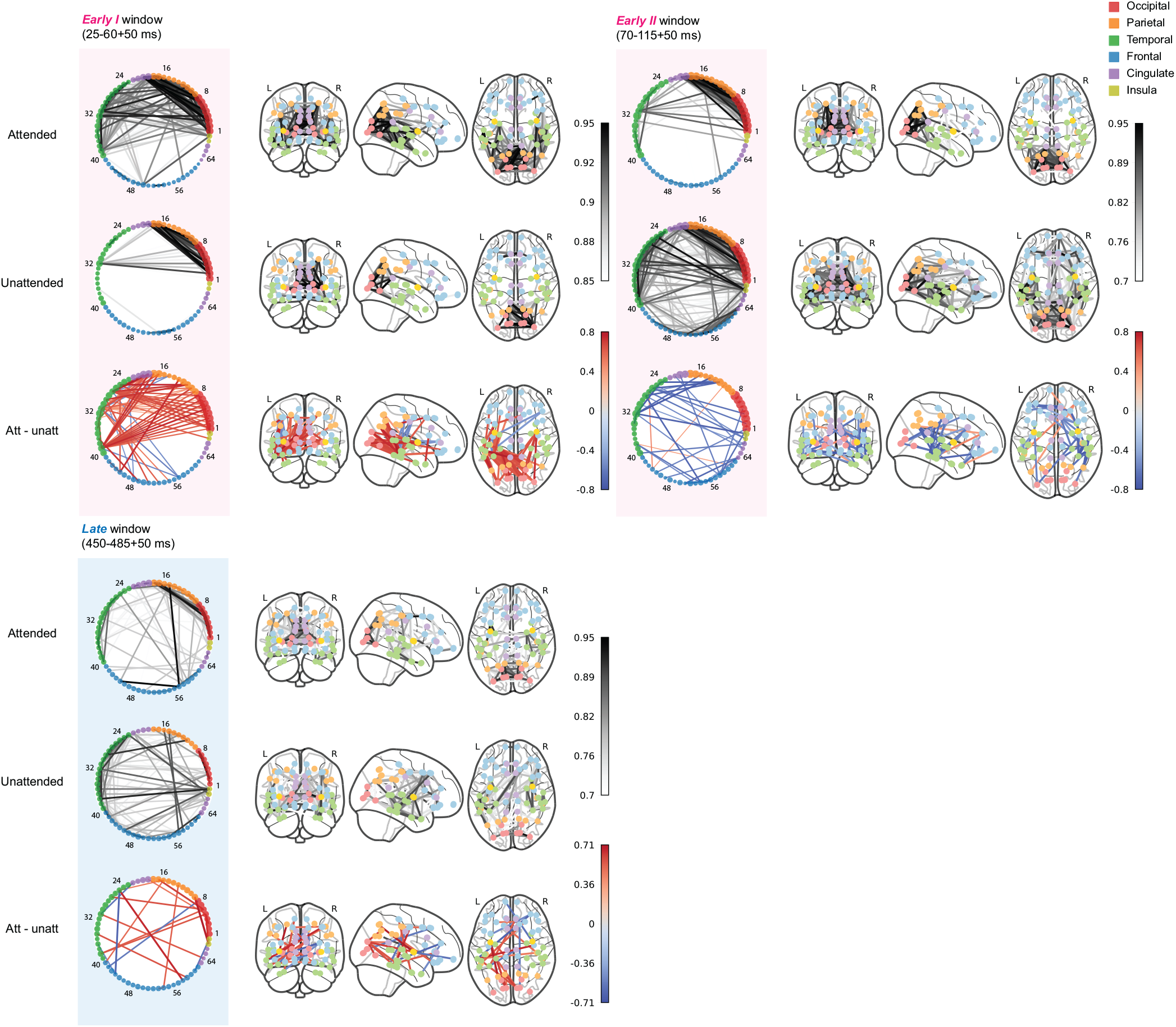
Visualization for averaged dynamic informational connectivity networks for T2 at different critical time windows. For each time window, the network is shown as a chord diagram in which each node represents an ROI; corresponding coronal, sagittal, and axial views are shown from left to right. The visualization of target attended networks is on the first row, and target unattended networks are on the second row. The darker the edges between the nodes, the stronger the connectivity is between them. The third row shows the difference in informational connectivity between attended and unattended conditions. The presented edges are p *<* 0.01 compared to a shuffled null distribution. Edges with higher values are shown in red, indicating greater connectivity during the attended than the unattended condition.

